# Formalizing metabolic-regulatory networks by hybrid automata

**DOI:** 10.1101/586198

**Authors:** Lin Liu, Alexander Bockmayr

## Abstract

Computational approaches in systems biology have become a powerful tool for understanding the fundamental mechanisms of cellular metabolism and regulation. However, the interplay between the regulatory and the metabolic system is still poorly understood. In particular, there is a need for formal mathematical frameworks that allow analyzing metabolism together with dynamic enzyme resources and regulatory events. Here, we introduce a metabolic-regulatory network model (MRN) that allows integrating metabolism with transcriptional regulation, macromolecule production and enzyme resources. Using this model, we show that the dynamic interplay between these different cellular processes can be formalized by a hybrid automaton, combining continuous dynamics and discrete control.

## 1 Introduction

Computational approaches in systems biology have become a powerful tool for understanding the fundamental mechanisms of cellular metabolism and regulation. However, the interplay between the regulatory and the metabolic system is still poorly understood. In particular, there is a need for formal mathematical frameworks that allow analyzing metabolism together with dynamic enzyme resources and regulatory events.

Concerning integrated modeling of metabolism and regulation, there exist approaches such as regulatory flux balance analysis (rFBA) [1] and Flexflux [2] that combine Boolean or multi-valued logical rules for transcriptional regulation with a steady-state stoichiometric model of metabolism. These techniques iterate flux balance analysis (FBA) by splitting the growth phase into discrete time steps. At each time step, the updated regulatory states are imposed as bounds on the reaction fluxes while ignoring the costs for enzyme production. At a different level, there exist methods to predict metabolic resource allocation considering enzyme-catalytic relationships, either at steady-state (RBA [3], ME models [4]) or in a dynamic setting (deFBA [5], cFBA [6]). But, regulation is not included in these approaches. Besides Boolean logic and stoichiometric models, piecewise-linear differential equations [7, 8] and other types of hybrid systems [9, 10] have also been used to study the dynamics of metabolic-genetic networks. Most of these studies, however, merely consider metabolism and regulation, and do not combine these with macromolecule production and enzymatic relationships.

In the present work, we introduce a metabolic-regulatory network model (MRN) extending the self-replicator system proposed in [11]. Our modeling framework allows integrating metabolism with transcriptional regulation, macromolecule production, enzyme resources, and structural building blocks. Using this framework, we show that the dynamic interplay between cellular metabolism, macromolecule production and regulation can be formalized by a hybrid automaton, combining continuous dynamics and discrete control. In this formalization, all metabolite concentrations are represented by continuous variables. The discrete states of the system are composed of all gene expression states for the regulated proteins, which include regulatory proteins and regulated enzymes. In each discrete state, the continuous variables evolve according to a system of differential equations that is specific for this state. The guard conditions for the state transitions depend on the amounts of the molecular species and associated thresholds. To validate our approach, we present a hybrid automaton for a simplified model of the bacterial diauxic shift.

Our formalization makes it possible to apply hybrid system tools for analyzing metabolic-regulatory cellular processes. Compared to the approaches mentioned above, this will allow us including regulation, macromolecule production and enzyme resources into the prediction of the dynamics of cellular metabolism.

## 2 Constructing the metabolic-regulatory interaction network

In Fig. 1 we show the typical regulatory processes in a cell in response to external/internal signals [12]. Regulatory proteins **RP** first sense extra- or intracellular signals and transmit these to the gene expression machinery, which then alters the production of proteins, in particular enzymes **E**. Changes of enzyme amounts will affect the metabolite levels which in turn provide feedback as internal signals. Furthermore, via the process of breaking nutrients into energy and building blocks (catabolism), metabolism influences the level of signaling by providing the precursors of regulatory proteins.

**Fig. 1:**
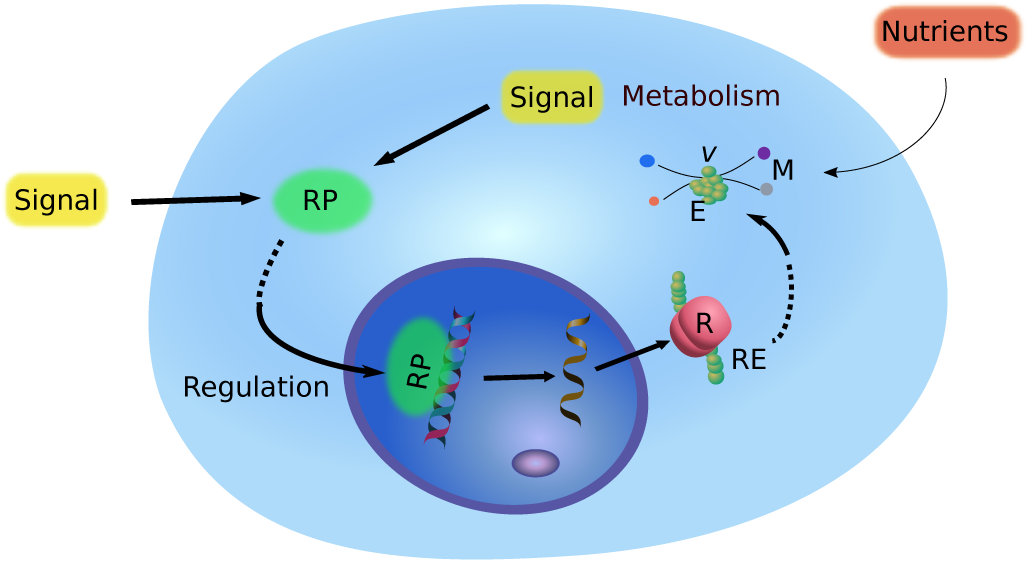
Typical metabolic-regulatory processes in response to external and internal signals.

We formalize the interactions between metabolism and regulation by a metabolic-regulatory network (MRN) that is given in Fig. 2. Regarding metabolism, **N** represents the set of external nutrients and **v**_**N**_ is the set of intermediate reactions that convert the nutrients into precursor metabolites **M**. The macromolecular production reactions **v**_**RP**_, **v**_**Q**_ and **v**_**E**_ = **v**_**RE**_ ∪ **v**_**NRE**_ use the precursors **M** to build regulatory proteins **RP**, non-catalytic macromolecules **Q**, and enzymes **E**. To keep the model simple, the set of enyzmes **E** contains all catalytic molecules, including transporters and ribosomes. However, we distinguish beween regulated enzymes **RE** and non-regulated enyzmes **NRE**, i.e., **E** = **RE** ∪ **NRE**. Non-catalytic macromolecules, termed as quota compounds **Q** [13], e.g. DNA and lipids, are included in the model because they are essential for growth and consume a lot of cellular resources.

**Fig. 2:**
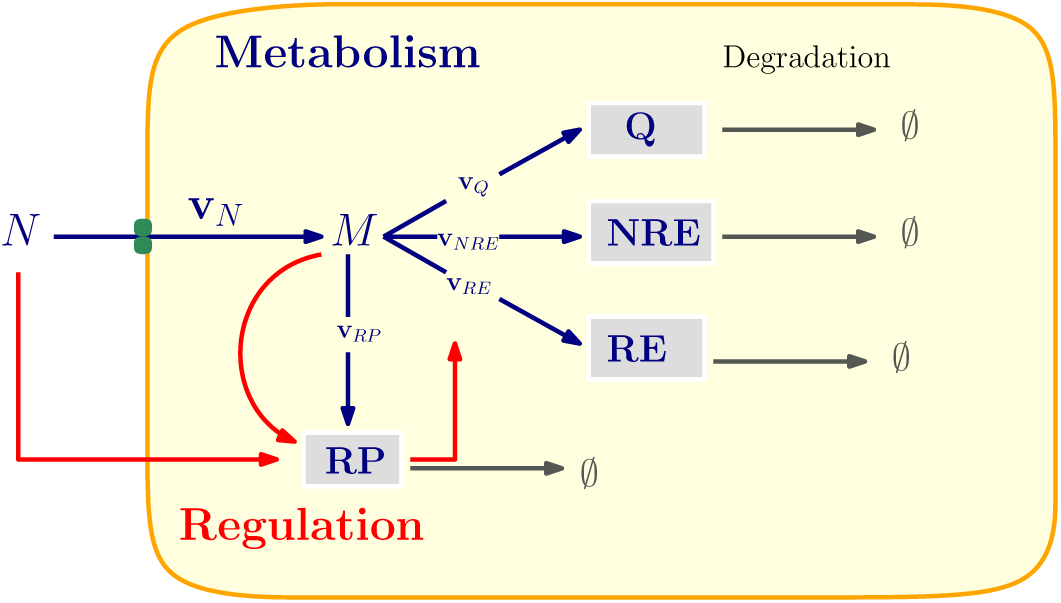
Schematic model of the metabolic-regulatory network (MRN). The metabolism is represented by the blue lines and the gene regulation by the red lines. Additionally, the degradation of macromolecules is shown by grey lines. We use **E** to represent all the catalytic macromolecules, which are classified as regulated enzymes **RE** and non-regulated enzymes **NRE**.

In the metabolic part, our network is inspired from the self-replication model by Molenaar et al. [11]. Compared to FBA-type models of metabolism these authors include metabolic resource allocation. However, they do not consider regulation. Similar to [1], we focus here on transcriptional regulation, i.e., we do not model post-transcriptional modifications.

In the following, we will show how the dynamics of the metabolic-regulatory model, i.e., the interactions between metabolism and regulation can be naturally described by a hybrid automaton.

## 3 Hybrid discrete-continuous dynamics

### 3.1 Continuous variables

Kinetic modeling of metabolic networks by ordinary differential equations (ODEs) has a long history in systems biology. Based on our metabolic-regulatory network (see Fig. 2), we define the set of molecular species

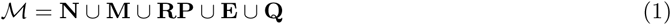

as the union of all nutrients, precursors, enzymes (including ribosome), regulatory proteins, and quota compounds. In a purely continuous modeling approach, the dynamics of the network would be described by a system of ODEs

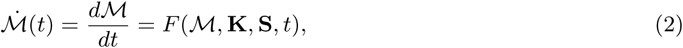

where **K** is the set of kinetic parameters, **S** is the stoichiometric matrix, and *t* denotes time. The function *F* represents the kinetic laws that govern the dynamics, which – depending on the molecular species – could be mass action, Michaelis-Menten, Hill kinetics etc.

### 3.2 Discrete states

Continuous modeling of gene regulatory networks is known to be very difficult due to the lack of the necessary kinetic data. Therefore, we adopt a more qualitative approach to include regulation in our model. It is based on the logical modeling framework pioneered in the 1970’s by L. Glass, S. Kauffman, R. Thomas et al., see [14] for a recent review. We assume that for each regulated protein *p* there are two possible states `on` and `off`, describing whether at a particular time *t* the gene encoding *p* is expressed or not.

Formally, for all *p* ∈ **RP** ∪ **RE**, we introduce a Boolean variable 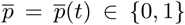 and a logical function *f*_*p*_: ℝ^*n*^ → {0, 1}. Here, the Boolean value 0 corresponds to `off` and the value 1 to on. Each function *f*_*p*_ is defined as a Boolean combination (using the Boolean operations *¬* (not), ∧ (and), ∨ (or)) of atomic formulae *x*_*i*_ ≥ *θ*_*i*_, where *x*_*i*_ is a real variable and *θ*_*i*_ is a constant. As an example, consider *f*: ℝ^2^ → {0, 1}, *f* (*x*_1_, *x*_2_) = (*x*_1_ ≥ 1) ∧ *¬*(*x*_2_ ≥ 2), for which we get *f* (1, 1) = 1 and *f* (1, 2) = 0. Overall, the regulation of our MRN is then formalized by a system of Boolean equations of the form

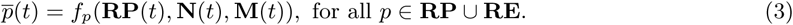

Here, *f*_*p*_ describes how the expression state of the gene encoding the regulated protein *p* depends on the current concentations of regulatory proteins, external nutrients, and intermediate metabolites.

### 3.3 Combining discrete and continuous dynamics in a hybrid automaton

Combining metabolism and regulation in this way leads to a hybrid discrete-continuous system. Here, all amounts of molecular species are modeled by continuous variables. However, the evolution of regulated proteins *p* is controlled by the expression state 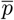 of the corresponding genes. Thus, depending on the discrete state 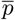, there are two different continuous dynamics. The system will jump from one discrete state to the other if some regulatory event occurs, see Fig. 3 for illustration.

**Fig. 3:**
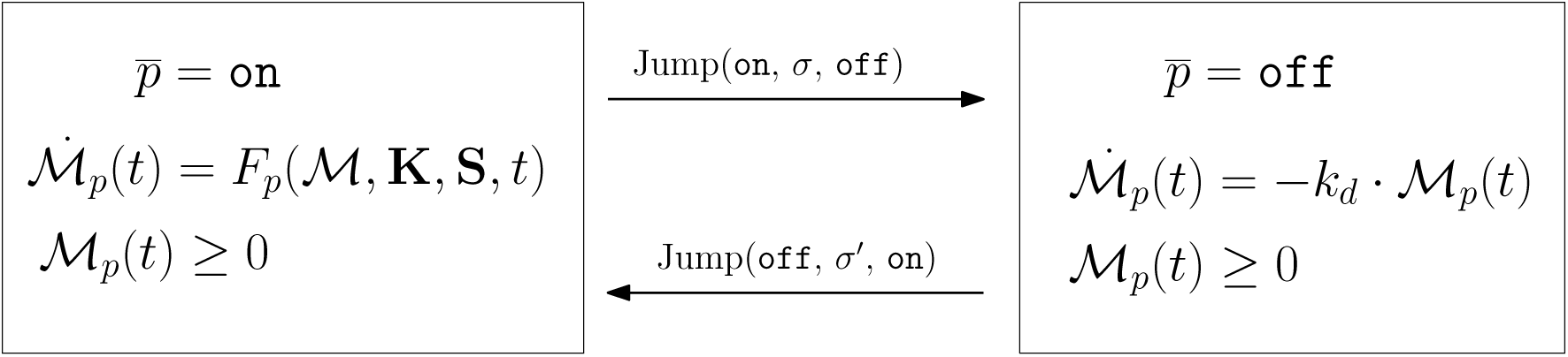
Graphical representation of the continuous evolution in the discrete states 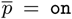 and 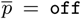, for regulated proteins *p* ∈ **RP** ∪ **RE**. In the on-state, protein production and degradation is described by the kinetic law *F*_*p*_(*·*), while in the `off`-state only degradation occurs, with kinetic constant *k*_*d*_.

Composing the discrete states together with their continuous dynamics for all regulated proteins *p* ∈ **RP** ∪ **RE** leads to a hybrid automaton

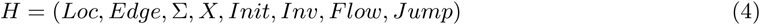

with the following components [15, 16]:

- *Loc* is a finite set of discrete states or locations. Here, *Loc* = {0, 1}^**RP**∪**RE**^ consists of all possible combinations of expression states 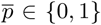 of regulated proteins *p* ∈ **RP** ∪ **RE**. In other words, for a MRN with *n* regulated proteins, there will be in total 2^*n*^ discrete states or locations in the hybrid automaton. However, not all of these have to be reachable from a given initial state.
- Σ is a finite set of events. In our case, these are given by the regulatory rules in Eqn. (3). For instance, a regulatory event *σ* can be that the amount of a regulatory protein *RP*_*i*_ is larger than a certain threshold *θ*_*i*_, which is expressed mathematically as *σ* = (*RP*_*i*_ ≥ *θ*_*i*_).
- *Edge* ⊆ *Loc* × Σ × *Loc* is the set of possible transitions from one location to another, which are labeled by an event from Σ.
- *X* is a finite set of real variables. In our case, *X* = ℳ= **N** ∪ **M** ∪ **RP** ∪ **E** ∪ **Q**. In each location *l*, these continuous variables evolve according to a specific dynamics depending on *l*, which is specified by the predicate *Flow*(*l*).
- *Init, Inv, Flow* are functions that assign logical predicates to each location *l* ∈ *Loc*:
  1. *Init*(*l*) is a predicate which describes the possible initial values for the continuous variables when the automaton starts its execution in state *l*.
  2. *Inv*(*l*) is a predicate which describes the possible values of the continuous variables when the control of the automata lies in *l*.
  3. *Flow*(*l*) is a predicate which describes the possible continuous evolutions when the control of the hybrid automaton is in *l*, for example by a system of ODEs.
- *Jump* is a function that assigns to each *e* ∈ *Edge* a predicate *Jump*(*e*) describing when the discrete change modeled by *e* is possible and what the possible updates of the continuous variables are when this change is made.

For a formal specification of the discrete-continous dynamics of the hybrid automaton *H* we refer to [15, 16]. In the next section, we explain the main principles by an illustrative example.

## 4 Biological application

Carbon catabolite repression (CCR) is a common phenomenon in bacteria, especially in *Escherichia coli* [17]. While these bacteria are able to grow on two different carbon sources, they do not consume these in parallel, but one after the other. To model the diauxic shift, we use the metabolic-regulatory network in Fig. 4, which was built based on the work in [1, 11]. In particular, we introduce a regulatory protein *RP* which, in presence of carbon source *C*_1_, inhibits the synthesis of transporter *T*_2_.

**Fig. 4:**
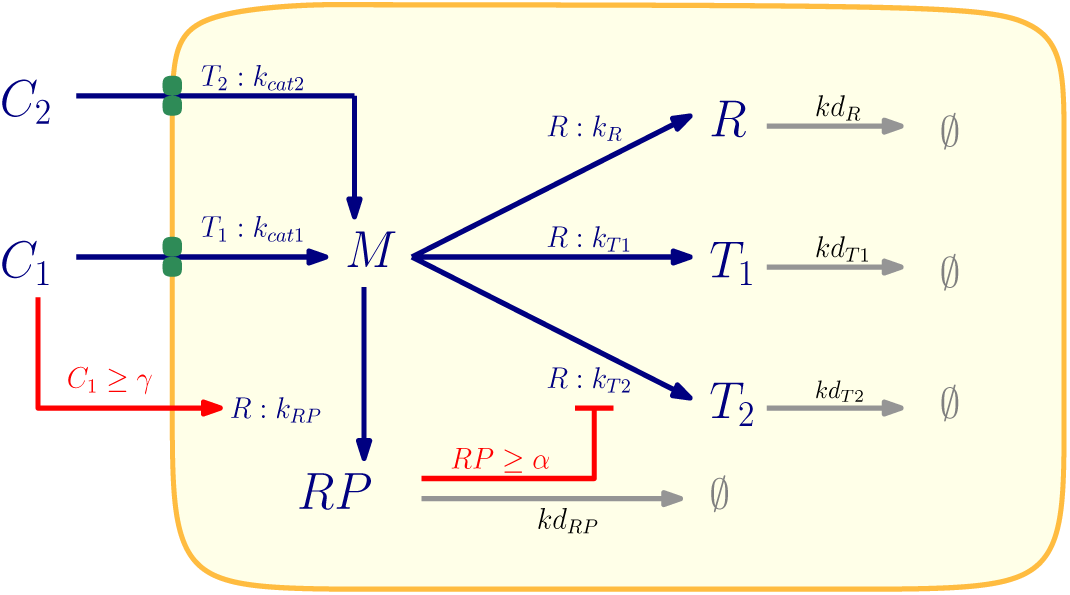
A self-replicator model with two regulatory rules. *C*_1_, *C*_2_ are the two carbon sources. *T*_1_, *T*_2_ are the enzymes for converting carbon sources into precursor *M*. *RP* is the regulatory protein. *R* represents the ribosome catalyzing the production of *RP, T*_1_, *T*_2_, *R* with turnover rate *k*_*cat*1_, *k*_*cat*2_, *k*_*R*_, *k*_*T* 1_, *k*_*T* 2_, *k*_*RP*_ and degradation rates are the corresponding turnover rates. *kd*_*R*_, *kd*_*T* 1_, *kd*_*T* 2_, *kd*_*RP*_. *γ, α* are the thresholds of the two regulatory rules.

### 4.1 Metabolic-regulatory network model of the diauxic shift

Starting from the generic model in Fig. 2, we consider two alternative carbon sources *C*_1_, *C*_2_ and a regulatory protein *RP* to build the MRN (see in Fig. 4). The two regulated proteins in our model are *RP* and *T* _2_, with corresponding Boolean variables 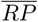 and 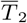. As shown by the red lines, the regulatory rules are the following: If the external amount of *C*_1_ is above the threshold γ, the gene encoding for *RP* is activated. If the amount of *RP* inside the cell is above the threshold *α*, the gene encoding for *T*_2_ is repressed. More formally:

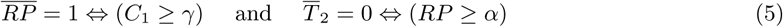

Together, these regulations ensure that *C*_1_ is the preferred carbon source for the model.

### 4.2 Hybrid automaton model of the diauxic shift

Next we construct the hybrid automaton *H*_*diaux*_ for the metabolic-regulatory network in Fig. 4. We get the continuous variables *X*_*diaux*_ = {*C*_1_, *C*_2_, *M, RP, T*_1_, *T*_2_, *R*} and the discrete locations 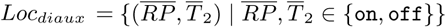. To specify the dynamics of *H*_*diaux*_, we use the graphical representation in Fig. 5, which is more intuitive than the formal definition according to Eqn. (4).

**Fig. 5:**
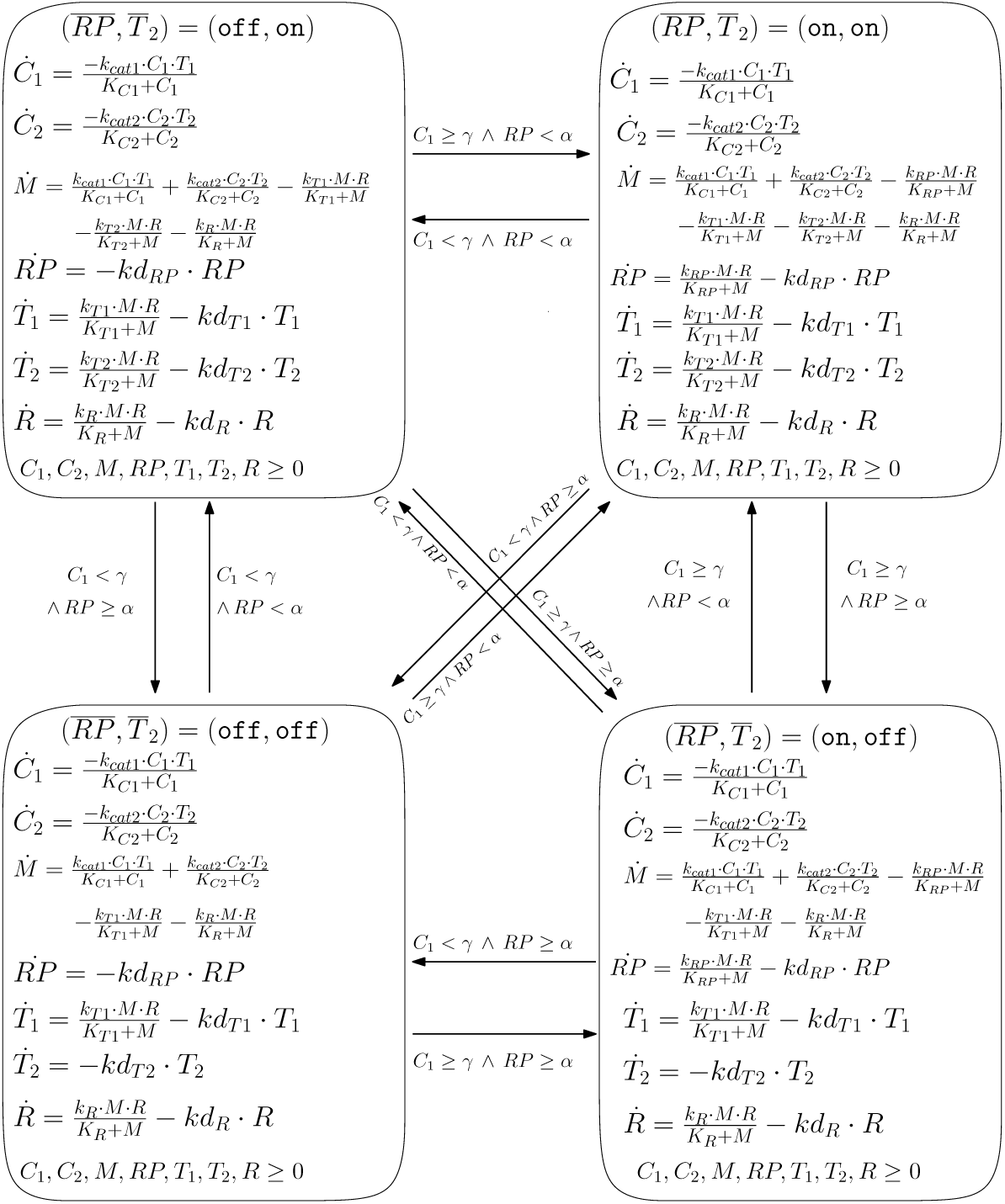
Graphical representation of the hybrid automaton *H*_*diaux*_.

The four nodes in Fig. 5 correspond to the discrete states or locations. Within each node, we specify the continuous dynamics of the molecular species by a set of ODEs and some invariants. The arcs represent the discrete jumps between different states. They are labeled by guards that the system has to satisfy in order to perform the corresponding transition.

If 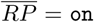, the regulatory protein *RP* is built from the precursor *M*, catalyzed by the ribosome *R*. Whenever *C*_1_ < *γ* is reached, 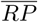 switches to `off` and *RP* is degraded with rate *kd*_*RP*_. Similarly, the enzyme *T*_2_ is produced if 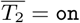, which is equivalent to *RP* < *α*, and degraded if 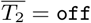. Due to mass balance, the dynamics of the precursor *M* also depends on the location and the regulatory control. The dynamics of the other variables *C*_1_, *C*_2_, *T*_1_, *R* is not directly controlled by the discrete states, but via the shared variables depends on them as well.

### 4.3 Exploring the dynamics of *H*_*diaux*_

Given the hybrid automaton *H*_*diaux*_, we explore the dynamics of the system for two different initial amounts of carbon source *C*_1_, see Fig. 6 and Fig. 7. In both cases, the kinetic parameters are the same (see Fig. 6) and the initial state is 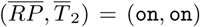. Following [18], we assume that the cell starts growing with some positive amounts of precursors and ribosomes, but without any enzymes. Thus, the initial enzyme amounts *T*_1_, *T*_2_ are set to 0, so that the model first has to produce them before carbon uptake can start. The only difference between the two simulations is that the initial value *C*_1_ is reduced from *C*_1_ = 30 in the first run to *C*_1_ = 3 in the second.

**Fig. 6:**
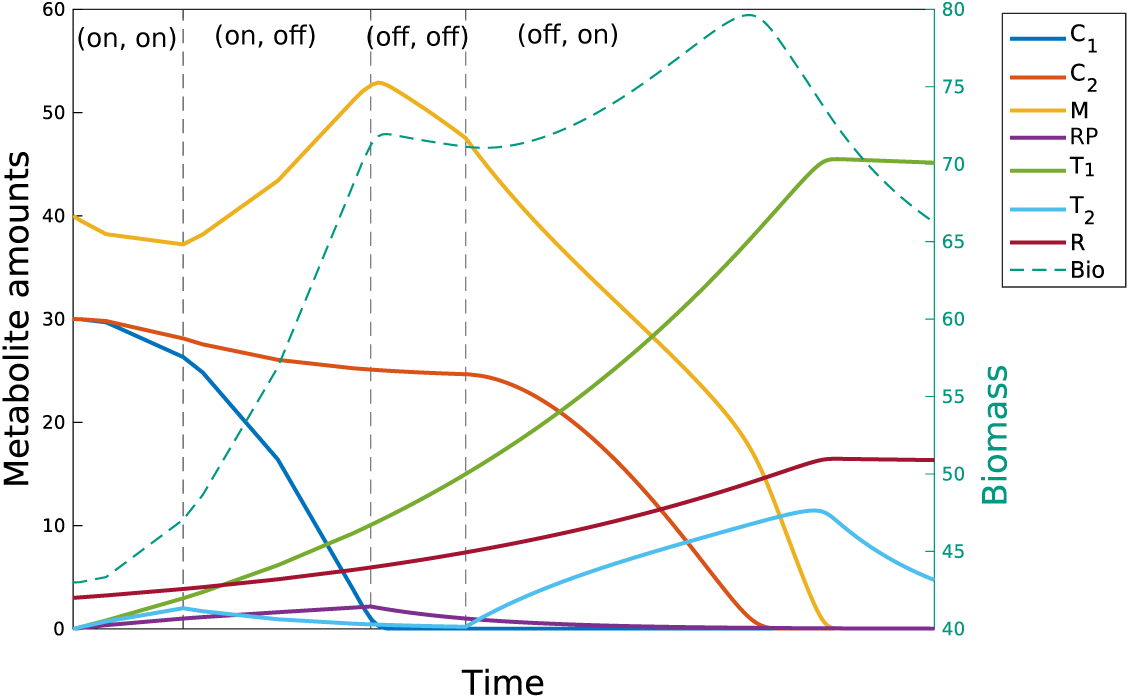
Simulation 1: Start in location 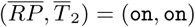 with initial values (*C*_1_, *C*_2_, *M, RP, T*_1_, *T*_2_, *R*) = (30, 30, 40, 0, 0, 0, 3) and kinetic parameters (*k*_*cat*1_, *k*_*cat*2_, *k*_*R*_, *k*_*T* 1_, *k*_*T* 2_, *k*_*RP*_, *kd*_*R*_, *kd*_*T* 1_, *kd*_*T* 2_, *kd*_*RP*_) = (0.3, 0.2, 0.03, 0.1, 0.1, 0.05, 0.001, 0.001, 0.1, 0.1). All Michaelis constants are assumed to be 1. Discrete state transitions are indicated by dashed lines, the thresholds are *α* = *γ* = 1.

**Fig. 7:**
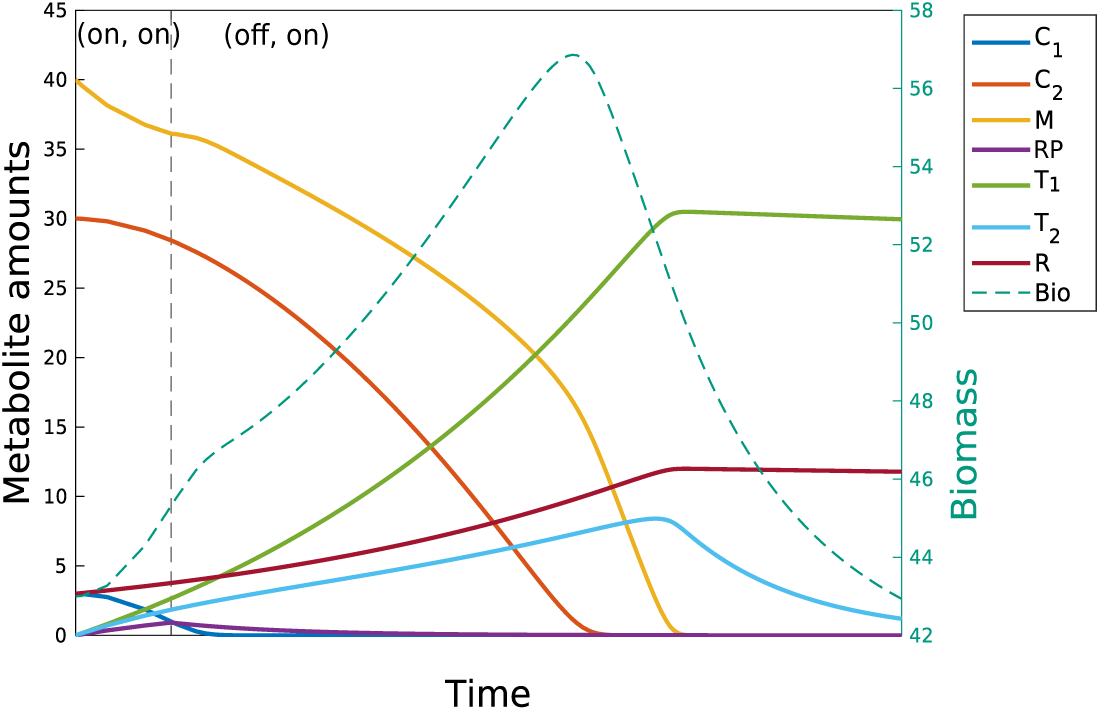
Simulation 2: Same data as in Simulation 1, except that initial value of *C*_1_ = 3. In this case, there is no lag phase. The model directly jumps from (`on, on`) to (`off, on`). since the initial amount of *C*_1_ is too small to inhibit the uptake of *C*_2_ via *RP*.

For Simulation 1, we can see in Fig. 6 that both *RP, T*_2_ initially increase because 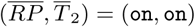. *C*_1_, *C*_2_ are consumed very slowly in the beginning, because the model is initialized with *T*_1_ = *T*_2_ = 0. When the amount of the regulatory protein *RP* reaches the threshold 1 and *C*_1_ is still larger than 1, 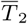 gets inactivated and the system will jump to the location 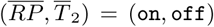. Now, the enzyme *T*_2_ stops being produced and with *T*_2_ getting close to 0, uptake of *C*_2_ is not possible anymore. Thus, the system can only use *C*_1_. The location 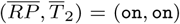 therefore indicates a first growth phase on the preferred carbon source *C*_1_. Once the guard condition *C*_1_ < 1, *RP* ≥ 1 is satisfied, the system will switch to the new state (`off`, `off`). This location represents the lag phase during diauxie, in which *C*_1_ is exhausted while the utilization of *C*_2_ is still repressed. Furthermore, the synthesis of the regulatory protein *RP* is inhibited and its amount decreases with the degradation rate *kd*_*RP*_. When the amount of *RP* reaches the threshold 1, 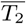 is turned on again and we get the final state (`off`, `on`). Now, the cell can produce *T*_2_ and will consume carbon source *C*_2_ until this is finally also exhausted.

In Simulation 2, the initial amount of *C*_1_ is reduced to 3. Starting again from the location (`on`, `on`), *RP* and *T*_2_ will increase and both carbon sources are assimilated. Due to the smaller initial value of *C*_1_ the event *C*_1_ < 1 ∧ *RP* < 1 is triggered first in this case. Hence the system directly jumps to (`off`, `on`) by skipping (`on`, `off`) and (`off`, `off`). As we can see in Fig. 7, there are no clear carbon switch and lag phase in this case. The regulatory protein *RP* does not reach the critical threshold *α* before *C*_1_ is depleted. From the biological viewpoint, this means that the amount of the preferred carbon *C*_1_ is too small to inhibit the uptake of *C*_2_.

Simulations 1 and 2 exhibit just two possible behaviors of *H*_*diaux*_. Fig. 5 shows all possible state transitions during diauxie based on our regulatory rules. Our results reveal that cellular behavior during diauxie is very sensitive to the initial conditions, which include the external environment and the intracellular status. This is consistent with the experimental results that the diauxie has high phenotypic heterogeneity [19, 20] and that the lag time is controlled by an inheritable epigenetic factor [21]. It also has been experimentally observed that the longer the cells grow in the preferred carbon source, the longer the lag phase is [22]. Our Simulation 2 shows there is even no lag phase when the initial amount of *C*_1_ is very small. The reason is still unclear. Yet our simulations suggest that when staying longer in the preferred carbon source, more regulatory proteins will be accumulated. Thus it will take more time to activate the second carbon source *C*_2_.

## 5 Conclusion

In this paper, we have proposed a new hybrid discrete-continuous modeling framework for metabolic-regulatory networks that integrates metabolism, transcriptional regulation, macromolecule production and enzyme resources. As a proof of principle, we have illustrated the approach on a small self-replicator model and applied it to study the diauxic switch in bacteria.

In our future work, we plan to construct larger and more realistic metabolic-regulatory networks and to apply existing simulation and analysis tools for hybrid systems in order to further study them. In particular, we plan to explore optimal control strategies for the hybrid automaton representing our MRN in continuation of our work on dynamic enzyme-cost flux balance analysis [5].

## 6 Acknowledgements

The authors would like to thank M. Köbis for his help in running the simulations. Lin Liu gratefully acknowledges support from the China Scholarship Council (CSC).

